# Antagonistic odor interactions in olfactory sensory neurons are widespread in freely breathing mice

**DOI:** 10.1101/847525

**Authors:** Joseph D. Zak, Gautam Reddy, Massimo Vergassola, Venkatesh N. Murthy

## Abstract

Odor landscapes contain complex blends of discrete molecules that each activate unique, overlapping populations of olfactory sensory neurons (OSNs). Despite the presence of hundreds of OSN subtypes in many animals, the overlapping nature of odor inputs may lead to saturation of neural responses at the early stages of stimulus encoding. Information loss due to saturation could be mitigated by normalizing mechanisms such as antagonism at the level of receptor-ligand interactions, whose existence and prevalence remains uncertain. By imaging OSN axon terminals in olfactory bulb glomeruli as well as OSN cell bodies within the olfactory epithelium in freely breathing mice, we found widespread antagonistic interactions in binary odor mixtures. In complex mixtures of up to 12 odorants, antagonistic interactions became stronger and more prevalent with increasing mixture complexity. Therefore, antagonism is a remarkably common feature of odor mixture encoding in olfactory sensory neurons and helps in normalizing activity to reduce saturation.

## Introduction

The number of distinct types of sensory neurons is usually far smaller than the number of distinct stimuli that an animal needs to detect, necessitating individual neurons to be receptive to multiple stimuli. This feature is especially relevant in the olfactory system where the number of odorous molecules vastly exceeds the repertoire of receptor types (Hopfield, 1999; Mainland et al., 2014; Su et al., 2009). Further, the olfactory world at any instant consists of complex mixtures of odorants, with individual olfactory receptors encountering multiple odorants at once. Therefore, how multiple ligands interact at single odorant receptors will define the mode of coding in, and the capacity of, the olfactory system. A large fraction of the olfactory sensory neurons (OSNs) in mammals express a family of odorant receptors that are G protein coupled receptors (Buck and Axel, 1991; Godfrey et al., 2004; Spehr and Munger, 2009). If the ligand-receptor binding kinetics are modeled with a single affinity parameter, simultaneous activation by multiple ligands will be simply determined by the relative affinities, until saturation (Arneodo et al., 2018; Dewan et al., 2018; Meister and Bonhoeffer, 2001; Wachowiak and Cohen, 2001). However, it is becoming increasingly clear that more complex interactions can occur.

Recent theoretical work has shown that a variety of non-linear interactions among multiple ligands at the same receptor can readily arise with a simple two-step model of receptor activation (Reddy et al., 2018). Experimental work *in vitro* in OSNs has suggested the existence of non-linear interactions, especially antagonism (Firestein and Shepherd, 1992; Kurahashi et al., 1994; Mathis et al., 2016; Rospars et al., 2008, 2000; Singh et al., 2019) (also bioRxiv preprints; Inagaki et al., 2019; Pfister et al., 2019; Xu et al., 2019). However, the prevalence of these interactions, especially in living animals within the constraints of natural sniffing dynamics, has not been explored. At a more basic level, evidence for multi-step receptor activation has also been sparse. For example, odorants with different affinities for a given receptor could also have distinct maximal activation if their efficacies are different (Rospars et al., 2008), but few studies have systematically explored this aspect.

Elucidating the principles of mixture interactions in OSNs is important for understanding odor coding. Odor identity and abundance are widely accepted to be represented by a combination of OSNs (Duchamp-Viret et al., 1999; Gottfried, 2010; Jinks and Laing, 1999). While each odorant activates a discrete pattern of sensory inputs, there can be considerable overlap in patterns of OSN activation corresponding to different odorants (Fletcher et al., 2009; Lin et al., 2006; Rubin and Katz, 2001; Shen et al., 2013). Since naturalistic odor stimuli are complex blends of many odorants, even relatively simple odor blends may saturate the entire complement of OSNs, thereby limiting their information coding capacity. More generally, even before saturation sets in, nonlinear interactions among multiple odorants in a given OSN may pose challenges for downstream decoding of odor identity (Howard and Gottfried, 2014; Wilson and Sullivan, 2011).

We have sought to systematically characterize mixture interactions at OSNs in their native environment using two-photon imaging of calcium responses to odor stimuli. Imaging populations of OSN axon terminals in the olfactory bulb glomerular layer afforded excellent signal-to-noise ratio and direct access to information conveyed to the brain. We then adapted a method to chronically image OSN somata in the olfactory epithelium of freely-breathing mice, which allowed us to bypass any influence of top-down neural feedback and to directly access the result of odor transduction. By delivering odors individually and in binary mixtures at concentrations that varied over three orders of magnitude, we were able to uncover evidence for multi-step activation of ORs and for widespread antagonistic interactions. We also found such antagonistic interactions in responses to complex mixtures containing up to 12 components in single OSNs. Our data strongly support a role for mixture-suppression of OSN activity as a normalizing mechanism in olfactory stimulus encoding.

## Results

### Antagonism measured in the glomerular layer of the olfactory bulb

Odor responses in OSNs can be measured with excellent sensitivity in the glomerular layer of the olfactory bulb, where axons of a given receptor type converge, allowing signal averaging. We used OMP-GCaMP3 mice and imaged axonal calcium responses through a cranial window over the dorsal surface of both olfactory bulbs. For two odorants, Methyl tiglate and Isobutyl propionate (Figure 1A), we measured odor responses in glomeruli across a range of concentrations spanning three orders of magnitude. We then made a binary mixture of the two odorants and measured OSN responses at the same glomeruli (Figure 1B-C). From nine mice, we identified 334 glomeruli that responded to either of the odors, or the mixture of the two. For this odor pair, many glomeruli reached different saturation levels for the same odor (Figure 1C). Across all glomeruli where saturation was achieved for a given odor, the distribution and mean Hill coefficients was similar (2.67 ± 0.16, n = 48 for Methyl tiglate and 2.53 ± 0.32, n = 22 for Isobutyl propionate). Early experiments with synthetic calcium indicators or intrinsic imaging of glomeruli estimated Hill coefficients to be close to 1, but more recent experiments appear to measure higher values (Arneodo et al., 2018; Dewan et al., 2018).

**Figure 1.**
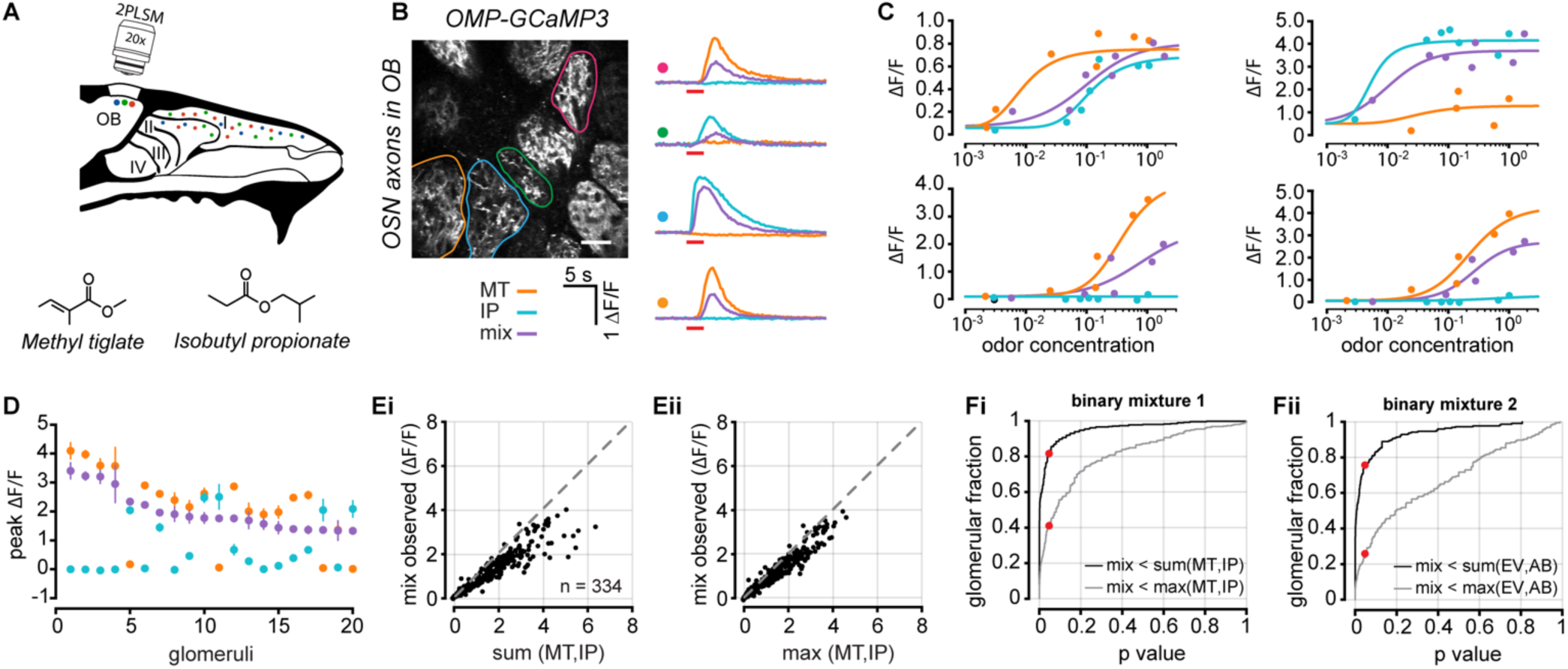
Optical measurements of antagonism measured in OSN axon terminals. **A**. Experimental setup. Odor responses in OSN axon terminals were measured at olfactory bulb glomeruli through a cranial window. **B**. Example image of glomeruli and selected ROIs. Odor responses for two odors (Methyl tiglate and Isobutyl propionate) and an equiproportionate mixture of both odors from selected glomeruli are shown on right as ΔF/F time courses. Colored circles correspond to ROIs in image. Red bar under traces denotes odor delivery time. Scale bar in image is 50 µm. **C**. Example dose response curves from four selected glomeruli. Each point is the average of 3-5 trials. Odor concentrations are normalized to the largest measured concentration (see Methods). **D**. Data from 20 randomly selected glomeruli. Each point is the mean of the three largest responses from each odor. Error bars are s.e.m. **E**. (**i**) Comparison of the observed mixture response against the linear sum of both mixture components or (**ii**) against the maximum response generated by either mixture component. **F**. Cumulative distribution of rank-sum p-values obtained for 334 glomeruli for linear sum comparison (black) and comparison to maximum component response (grey). Red dots mark p-values < 0.05. **G**. Same as *F*, but for an odor pair with highly overlapping responses (n = 226 glomeruli; extended data in Supplemental Figure 1).

To estimate antagonism at each glomerulus, we first calculated the average response to the three highest odor concentrations of each individual odor (Figure 1C-D). We then tested whether the observed mixture responses for the same three concentrations were significantly below the linear sum of the response to the mixture components (rank sum test of trial replicates). Significant mixture suppression was found in 80.5% (277/344) of glomeruli that responded to at least one odor (Figure 1Ei, Fi).

There are, however, two important considerations in interpreting our result. First, it remains possible that the largest calcium signals observed for individual odors may represent the average maximum physiologically-bounded firing rate across all OSNs of a common type. Similarly, the largest signals we observed for individual odors may be bounded by calcium indicator saturation. Given these constraints, it is plausible that the observed mixture responses can never reach the linear sum of the two mixture components and are simply bounded by the largest response observed for either component alone. To account for this possibility, we compared between the observed mixture responses and maximum response for either odor. When using this more conservative metric, we found evidence for antagonism in 41.7% (144/344) of the glomeruli (Figure 1Eii,Fi). These data indicate that, even by a conservative measure, antagonism is remarkably prevalent in OSNs, when estimated from glomerular activity.

In these experiments, glomerular responses were highly non-overlapping for the odor pair tested. To ensure that antagonism did not arise from unique interactions between these two odors, and to demonstrate generalization of the phenomenon with other odor pairs, we repeated our experiments using another odor pair, Ethyl valerate and Allyl butyrate. This pair was specifically selected because of their highly overlapping OSN activation (Supplemental Figure 1). Mixture interactions for this pair of odors were in close agreement with those estimated above for the other odor pair. From 226 glomeruli, we found antagonism in 76.6% (173/226) of glomeruli when comparing the sum responses and 25.7% (58/226) when comparing against the maximum (Figure 1Eii). Similar to our previous odor pair we found that the distribution of Hill coefficients was remarkably similar (2.36 ± 0.19, n = 44 for Ethyl valerate and 2.29 ± 0.21, n = 39 for Allyl butyrate).

### Imaging odor responses in individual OSNs

Our glomerular imaging results suggest that mixture-suppression through antagonism is widespread. However, an alternative mechanism that could produce such an effect may operate through GABAb- or D2-mediated suppression of OSN axon terminals (Aroniadou-Anderjaska et al., 2000; Ennis et al., 2001; Fleischmann et al., 2008; McGann et al., 2005).

To avoid circuit interactions between different OSNs, we directly imaged OSN somata *in situ* in the olfactory epithelium (Iwata et al., 2017), where there is no evidence for efferent modulation that is directly linked to olfactory circuitry (Lucero, 2013). We first characterized the odor turning and response dynamics of individual cells to determine whether the response properties of sensory neurons in the dorsal recess (zone 1) of the olfactory epithelium (Kobayakawa et al., 2007; Ressler et al., 1993) are congruent with those observed in glomeruli in the dorsal olfactory bulb, and to ensure that we sample from a heterogeneous population of OSNs.

We used a panel of 32 monomolecular odors (see Methods) that activate glomeruli on the dorsal olfactory bulb across a wide range of densities (Figure 2D,F). We then imaged individual cells in the epithelium (Figure 2B) and compared their tuning to glomeruli in the olfactory bulb. Across odors, there was a strong relationship between the fraction of activated glomeruli and individual OSNs (R^2^ = 0.562, P < 0.001, Figure 2G). We found a similar relationship when we compared the mean response magnitude at both imaging sites across all glomeruli and somata (R^2^ = 0.375, P < 0.001, Figure 2I). These data provide new functional evidence that support anatomical data for a conserved zonal organization between the olfactory epithelium and the olfactory bulb (Kobayakawa et al., 2007).

**Figure 2.**
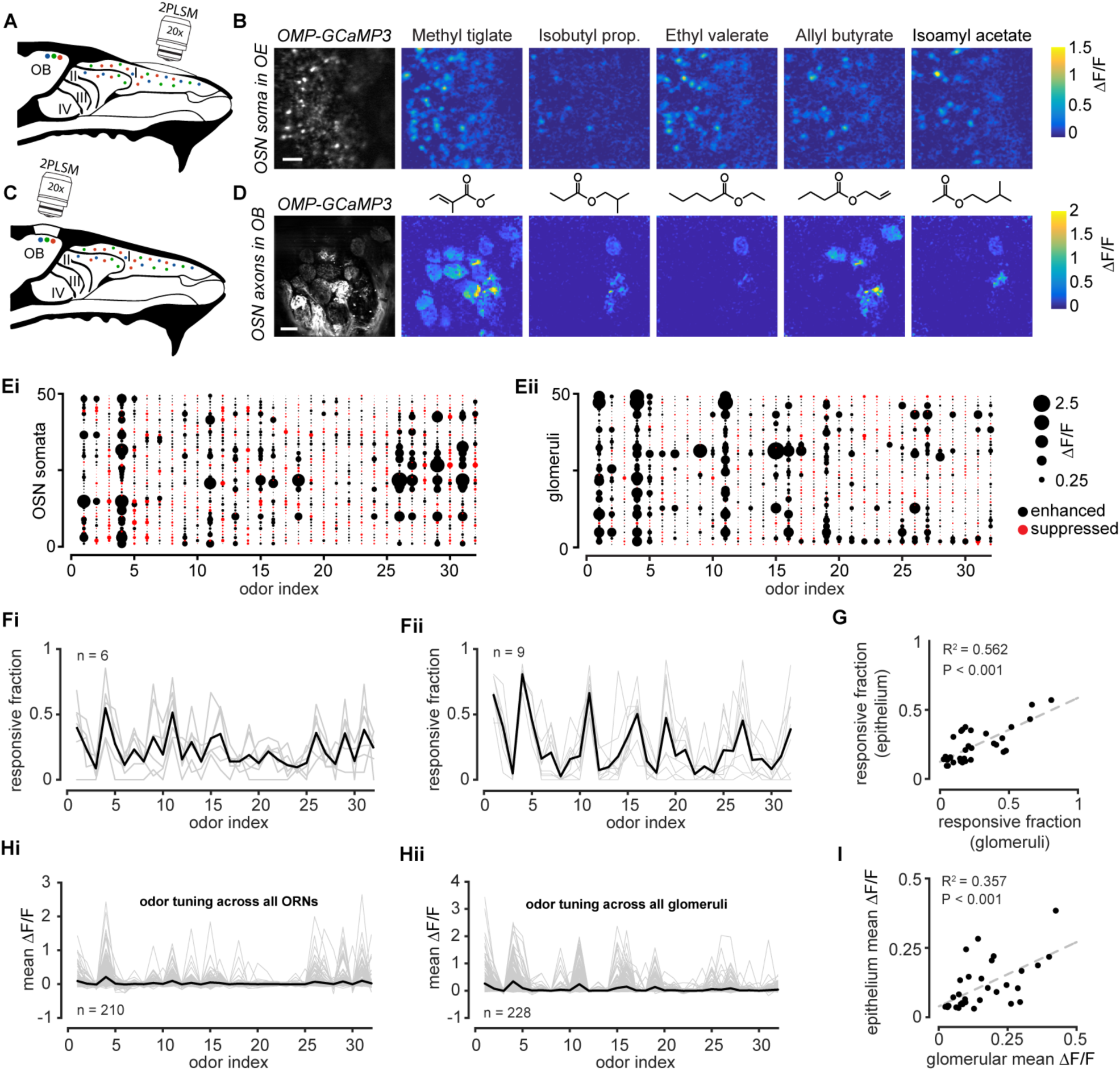
Odor tuning in the dorsal recess of the olfactory epithelium is similar to the dorsal olfactory bulb. **A**. Experimental setup for imaging OSN somata in the epithelium. **B**. Resting fluorescence in the soma of GCaMP3-expressing OSNs in the olfactory epithelium (OE) and heatmaps of selected odor responses. Scale bar is 30 µm. **C**. Experimental setup for imaging axon terminals of OSNs in the glomerular layer of the olfactory bulb **D**. Resting fluorescence of glomeruli and heatmaps of selected odor responses. Scale bar is 100 µm. **E**. Odor tuning of (**i**) 50 randomly selected OSNs (**ii**) and 50 glomeruli across 32 monomolecular odors. **F**. Fraction of (**i**) somata or (**ii**) glomeruli that responded to each of the odors in a given imaging field. Grey lines are individual fields and dark line is the mean across all imaging fields. **G**. Scatter plot of the responsive fraction across all somata and glomeruli for each of the 32 odors used in our panel. Dashed line is the linear regression to the data (R^2^ = 0.562, P < 0.001). **H**. Odor tuning of (**i**) 210 OSN somata and (**ii**) 228 glomeruli. Grey lines are individual somata or glomeruli and dark line is the mean across all ROIs. **I**. Scatter plot of the mean response across all somata and glomeruli for each of the 32 odors used in our panel. Dashed line is the linear regression to the data (R^2^ = 0.357, P < 0.001).

We also found that the calcium response kinetics across all OSNs in a field of view are highly diverse, and that the response waveform for individual cells is remarkably stable across trials of the same odor when respiration is stabilized (Supplemental Figure 2). Together these data suggest that a diverse and heterogeneous array of receptor subtypes are present within a relatively restricted patch of olfactory epithelium.

### Antagonism in individual sensory neurons

Given that OSNs in the dorsal recess share many of the same odor tuning characteristics as dorsal olfactory bulb glomeruli, we tested whether antagonism could be observed at the single cell level using the odor pair we used for imaging glomerular responses. We repeated the experiment described in Figure 1 and collected data from 964 individual sensory neurons using the odor pair, Methyl tiglate and Isobutyl propionate. We again detected non-linear interactions between the odors Figure 3B-C. Mixture-suppression was readily observed in individual cells (54.6% (527/964) for sum comparison and 22.0% (212/964) maximum comparison; Figure 3E,Fi). The fraction of cells that showed mixture-suppression was smaller than in the glomerular data, which may be attributed to noise from two sources. First, the measured signal at glomeruli represents the population response across all ~10,000 OSNs of a common subtype, compared to single OSN cell bodies in the epithelium imaging. The second source of noise may arise from image acquisition, through the number of pixels contributing to the measured signal. While we typically imaged 10-20 densely packed glomeruli per field of view, in the epithelium, we imaged dozens to hundreds of sparsely arranged cells, thereby substantially reducing the number of pixels contributing to our signal.

**Figure 3.**
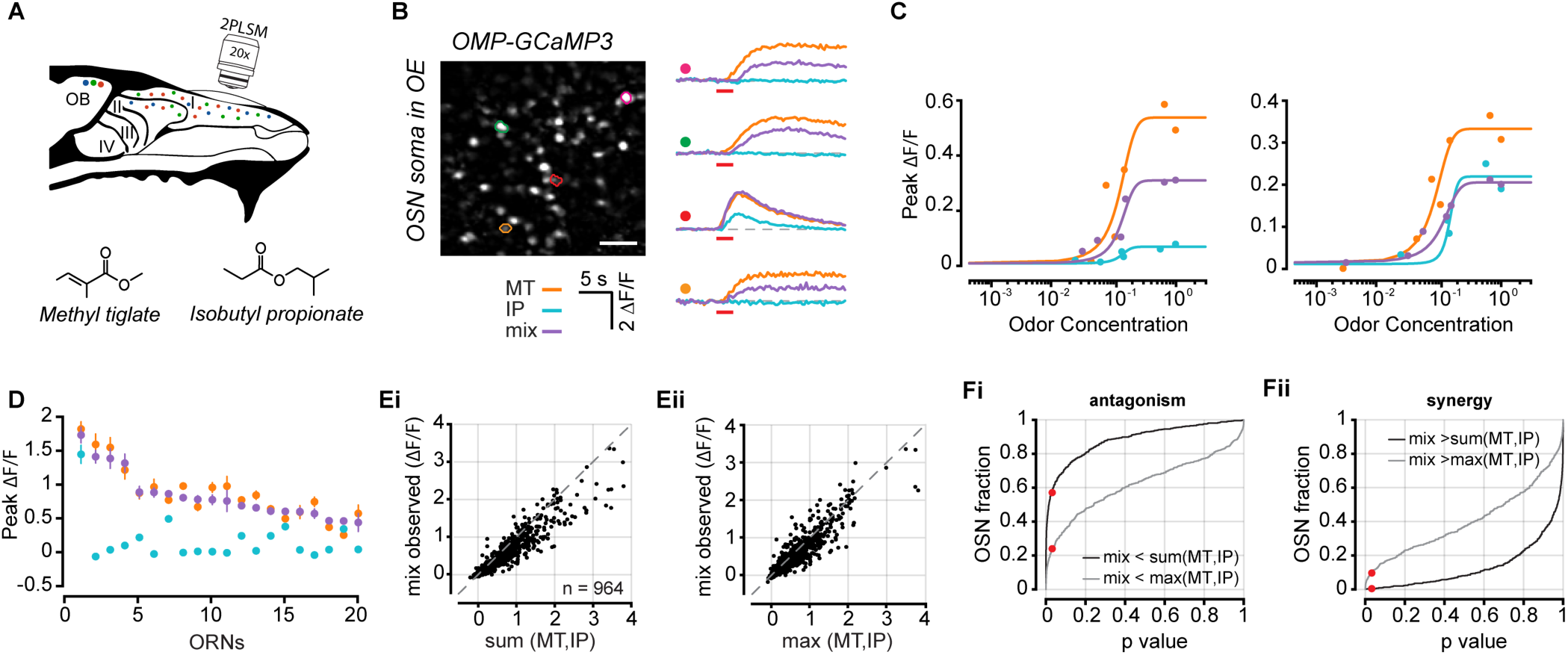
Antagonism in individual sensory neurons. **A**. Experimental setup for imaging OSN somata in the epithelium. **B**. Example image of OSN somata and selected ROIs. Odor responses for two odors (Methyl tiglate and Isobutyl propionate) and an equiproportionate mixture of both odors from selected OSNs are shown on right as ΔF/F time courses. Red bar under traces denotes odor delivery time. Scale bar in image is 30 µm. **C**. Example dose response curves from two selected OSNs. Each point is the average of 3-5 trials. **D**. Data from 20 randomly selected OSNs. Each point is the mean response of the three highest odor concentrations for each odor. Error bars are s.e.m. **E**. (**i**) Comparison of the observed mixture response against the linear sum of both mixture components or (**ii**) against the maximum response generated by either mixture component. **F**. (**i**) Cumulative distribution of rank-sum p-values obtained for 964 OSNs for linear sum comparison (black) and comparison to maximum component response (grey). Red dots mark p-values < 0.05. (**ii**) Cumulative distribution of left-sided rank-sum p-values for linear sum comparison (black) and comparison to maximum component response (grey) to identify synergistic mixture interactions.

To ensure that our observations did not arise from more noisy measurements at the single-cell level, we measured the fraction of mixture responses that exceeded the linear prediction of the mixture components, a phenomenon known as synergy. We first measured the fraction of mixture responses that exceeded maximum of either of the mixture components. We found only 11.8% (114/964) of OSNs exceeded the maximum response of either component (Figure 3E,Fii); however, it should be noted that for synergy to occur the measured OSN response should exceed the linear summation of the response of both odor components. In our dataset, we found only 5/964 cells where the mixture response was significantly greater than the sum of the mixture components. Both fractions are well below those we measure for antagonism and consistent with previous reports that antagonism is the predominant mixture interaction when compared with synergy (Duchamp-Viret et al., 2003).

### Antagonism in complex odor blends

After establishing that antagonism is a prevalent feature of odor mixture encoding in OSNs, we next wanted to understand the relationship between the complexity of an odor blend (that is, the number of elements in the mixture) and non-linearities in OSN responses. We imaged single OSN responses to 16 monomolecular odors as well as random blends of 2, 4, 8, or 12 odors from this panel (Methods; Figure 4C). From the OSN responses to each of the single odors, we then made linear predictions for each of the mixture complexities.

**Figure 4.**
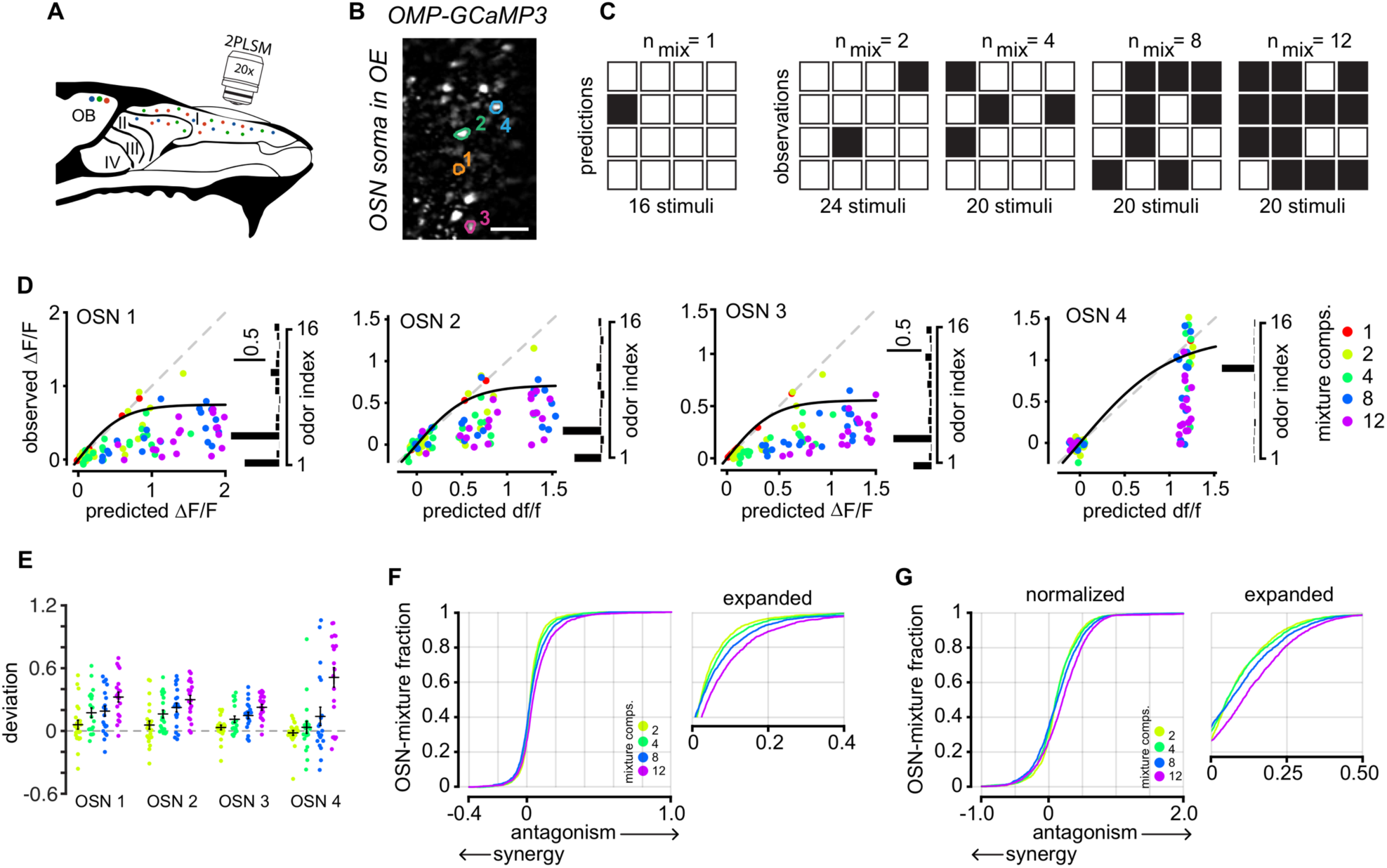
Antagonism through complex mixtures in single neurons. **A**. Experimental setup for imaging OSN somata in the epithelium. **B**. Example image of OSN somata and selected ROIs. Scale bar in image is 20 µm. **C**. Stimulus design for complex mixtures. OSN responses to each of 16 components are measured. Odor mixtures are made from subsets of 2, 4, 8, or 12 of the 16 individual odor components. **D**. Example mixture responses from the OSNs outlined in *B*. Each point is the average of three trials. The data from each OSN is fit with a sigmoid that reflects the maximum response from that cell. Odor tuning profile for each OSN is shown on the right. **E**. Deviation from each mixture response to the sigmoidal fit the data. Positive values reflect antagonism and negative values reflect synergy. Mean of all mixtures of a given complexity is the horizontal black bar and error is s.e.m. **F**. Cumulative distribution of all deviations from the sigmoidal fit for each mixture complexity. Data collected from 129 OSNs in 3 mice. Expanded traces are on the right. **G**. Data were normalized for each OSN by dividing each mixture response by the asymptote of the sigmoidal fit to the data. Antagonism is not overrepresented in a small number of strongly responding cells.

For all mixture complexities, we routinely observed OSN responses that were far less than the linear prediction made by the summation of the individual mixture components (Figure 4D). However, OSNs may never achieve the response predicted by linear summation, due to either firing rate or indicator saturation. To account for this possibility, for each OSN, we fit the data across all mixture complexities with a sigmoid with an initial slope of 1 and reaching an asymptote at the top 0.10 quantile of all observed mixture responses (n = 100). For each cell, we then calculated the deviation of each mixture response to the fit of the data. We observed non-linear mixture responses in OSNs at all mixture complexities, although both the frequency and magnitude of such interactions increased with mixture complexity (Figure 4F, n = 1800, n = 1596, n = 1861, n = 1948 OSN-odor mixture pairs, for 2-, 4-, 8-, and 12-part mixtures respectively). For this analysis, we only considered OSN-mixture pairs where the predicted OSN response was > 0 ΔF/F. Importantly, mixture suppression was not the result of large trial-to-trial variability where a single trial strongly influenced the mean as the error for each mixture complexity was well below the sigmoidal fit to the data (Supplemental Figure 3).

To exclude the possibility that a small number of cells most strongly contribute to the antagonism we observe, we normalized all measurements from each cell to the asymptote of the sigmoidal fit to the data and repeated our analysis. Here, we again observe that antagonism increases with mixture complexity (Figure 4G). As a further confirmation of our finding, we repeated this experiment in OSN axon terminals in the glomerular layer. We collected data from 39 glomeruli in 4 mice. We again observed strongly non-linear mixture interactions that increased with mixture complexity (Supplemental Figure 3; n = 673, n = 637, n = 733, n = 750 glomeruli-odor mixture pairs, for 2, 4, 8, and 12-part mixtures respectively). Together, our results demonstrate that input to the olfactory system may be normalized through mixture suppression of OSN activity.

## Discussion

It has been known for some time that individual odorant receptors can respond to multiple odors (Malnic et al., 1999), so a reasonable question is how these receptors will respond to natural stimuli, which are almost always mixtures of odors. Odor mixture interactions have been described behaviorally for many decades (Bell et al., 1987; Frank et al., 2017; Jinks and Laing, 2001). One form of interaction is where binary (and larger) mixtures tend to have reduced perceived intensity (“hypoadditivity”). Odors can also be masked or harder to identify in mixtures, which could be due to reduced sensation of specific components. The origin of this perceptual odor suppression in mixtures is uncertain. Some of it could arise at the periphery, including at the receptors themselves. We recently proposed a simple model based on the two-step activation of odorant receptors, and noted that even a modest decorrelation of binding affinity and activation efficacy across odors could lead to antagonistic interactions, which will lead to mixture suppression (Reddy et al., 2018).

Here, we have used direct imaging of OSN somata, as well as more conventional glomerular imaging, to show that mixture suppression (by inference antagonistic interactions) is widespread. We adapted a method first reported by Imai and colleagues (Iwata et al., 2017), to directly image the somata of OSNs *in vivo*, in freely breathing mice. We obtained responses of OSNs to a diverse array of odorants and found that the statistics of population responses were similar to that obtained for glomeruli. Interestingly, inhibitory responses were more easily detected in glomeruli than in OSN cell bodies, which might reflect glomerular averaging of heterogeneous responses of OSNs (Grosmaitre et al., 2006). This OSN imaging method is likely to be valuable for understanding the fundamental properties of sensory coding and plasticity at the periphery, especially with chronic imaging.

Previous studies using binary mixture stimuli have offered conflicting evidence for nonlinear interactions (Fletcher, 2011; Grossman et al., 2008; Lin et al., 2006; Rospars et al., 2008). Evidence for relatively linear summation of odor responses to mixtures has been offered by studies that have imaged glomerular responses at low spatial and temporal resolution (Fletcher, 2011; Grossman et al., 2008; Lin et al., 2005), but these studies did not systematically vary the concentration of odorants. Extracellular spike recordings from single rat olfactory sensory neurons have indicated that a simple competitive binding model can explain the responses in about half the cases (Rospars et al., 2008). In the rest of the cases, more complex interactions including antagonism (competitive or non-competitive) and masking were needed to explain the observations (Rospars et al., 2008). Insect olfactory receptors also exhibit such nonlinear interactions (Silbering and Galizia, 2007; Su et al., 2011). Recently, we offered a principled explanation for all these different effects using a model that decouples the binding affinity of odorants to receptors and their activation efficiency (Reddy et al., 2018). Then, depending on the statistics of these properties in the repertoire of odorants encountered by an animal, the population of odorant receptors could exhibit varying degrees of antagonism or synergy.

In this work, we used binary mixtures and concentration variation over three orders of magnitude, to find that suppressive mixture interactions were remarkably widespread. We first demonstrated mixture suppression with glomerular imaging. Since glomerular responses are averages of many hundreds of OSN axons, they will overlook any heterogeneity. Glomerular responses may also be influenced by lateral interactions within the olfactory bulb networks, particularly due to GABAb-mediated presynaptic inhibition that can alter OSN calcium signals (Aroniadou-Anderjaska et al., 2000; Wachowiak et al., 2005). However, there is strong evidence that GABAb-mediated reduction of OSN responses are largely intraglomerular (McGann et al., 2005), which will simply serve as an automatic gain control, hence insensitive to the identity of the odorant. As an additional check, we chose two distinct odor mixtures with differing extents of overlap in glomerular activation. We found very similar mixture effects for both pairs of odorants, which indicates that lateral interactions among glomeruli are unlikely to affect our conclusions.

To circumvent circuit and feedback interactions, we imaged OSNs directly and found that binary mixture suppression was just as widespread as observed with glomerular imaging. Since these interactions were observed at the earliest stages of odor encoding, in OSNs, the most likely source of mixture interactions is odorant receptors themselves. Antagonistic interactions have been hinted at previously, but we find direct evidence for widespread occurrence of this phenomenon. Going beyond binary mixtures, we used odor blends of increasing complexity, up to 12 components, to ask how mixture responses compared to the expected linear summation of individual responses (below saturation). This allowed us to ask how OSNs respond to more complex mixtures. We found that the degree of mixture suppression was greater with increasing number of components, which is expected from statistical consideration of antagonistic interactions (Reddy et al., 2018). This feature can be rationalized as increasing normalization of population responses, which we have shown previously to allow greater information transfer about odor identity, when saturation threatens degradation of information (Reddy et al., 2018).

Beyond our theoretical observations, there are other more recent experimental studies (in bioRxiv preprints) that report related effects (Inagaki et al., 2019; Pfister et al., 2019; Xu et al., 2019). Our work is unique in the following regard: (1) we report responses in OSNs in intact, freely-breathing mice, responding to vapor phase odorants inhaled physiologically, (2) we report fast, real-time responses with a variety of odorants and using both OSN and glomerular imaging and (3) we report responses to mixtures of high complexity (up to 12 odors), well beyond just binary mixtures.

Given this widespread existence of normalization, what advantage would a similar phenomenon at the level of sensory transduction itself confer to the system? Having normalization occur early in the sensory hierarchy helps avoid saturation early on and preserve more information about the stimulus to be conveyed downstream. Indeed, in our previous work, we demonstrated using information theoretic calculations that a target odorant embedded in a complex mixture can be more easily detected with antagonistic interactions that lead to sparser representation (Reddy et al., 2018). The improved performance with antagonism exists for a wide range of receptor “tuning widths” (i.e., the average number of activated glomeruli per single odorant). Importantly, normalization at the level of receptors leading to sparser, more informative representation, comes for free without additional circuit burden.

## Methods

### Experimental model and subject details

Adult heterozygous OMP-GCaMP3 mice (Isogai et al., 2011) of both sexes were used in this study. All animals were produced from a breeding stock maintained within Harvard University’s Biological Research Infrastructure. All animals were between 20 and 30 g before surgery and singly housed following any surgical procedure. Animals were two to six months old at the time of the experiments. All mice used in this study were housed in an inverted 12-hour light cycle and fed *ad libitum*. All the experiments were performed in accordance with the guidelines set by the National Institutes of Health and approved by the Institutional Animal Care and Use Committee at Harvard University.

### Olfactory bulb craniotomy

A craniotomy was performed to provide optical access to both olfactory bulbs as previously described (Zak et al., 2018). Animals were allowed to recover for at least three days. Prior to each imaging session, animals were anesthetized with a mixture of ketamine and xylazine (90% of dose used for surgery) and body temperature was maintained at 37 °C by a heating pad. Respiration was measured through an airflow sensor (Honeywell) (Bolding and Franks, 2017) during most experiments and maintained between 0.5 and 1.5 Hz (traces in Supplemental Figure 2F).

### Olfactory epithelium thinned skull procedure

Mice were anesthetized using the same procedure and all pre-surgical methods through head plate implantation are the same as the craniotomy. The cranial bones over the olfactory epithelium were thinned with a dental drill and blade until transparent(Iwata et al., 2017). The thinned area of skull was then covered with cyanoacrylate adhesive (Loctite) and a class coverslip was implanted in the adhesive. Dental cement was then used to form a well over the thinned section of skull.

### Multiphoton Imaging

A custom-built two-photon microscope was used for *in vivo* imaging. Fluorophores were excited and imaged with a water immersion objective (20X, 0.95 NA, Olympus) at 920 nm using a Ti:Sapphire laser (Mai Tai HP, Spectra-Physics). Images were acquired at 16-bit resolution and 4-8 frames/s. The pixel size was 0.6 μm OSN somata imaging and 1.2-2.4 μm for imaging glomeruli. Fields of view ranged from 180 × 180 μm in the epithelium to 720 x 720 μm in the glomerular layer. The point-spread function of the microscope was measured to be 0.51 × 0.48 × 2.12 μm. Image acquisition and scanning were controlled by custom-written software in LabView (National Instruments). Emitted light was routed through two dichroic mirrors (680dcxr, Chroma and FF555-Di02, Semrock) and collected by a photomultiplier tube (R3896, Hamamatsu) using filters in the 500–550 nm range (FF01–525/50, Semrock).

### Odor stimulation

Monomolecular odorants (Sigma or Penta Manufacturing) were used as stimuli and delivered by custom-built 16 channel olfactometers controlled by custom-written software in LabView (Zak et al., 2018). For binary mixture experiments, the initial concentration series for each odor was between 0.08% - 80% (v/v) in mineral oil and further diluted 16 times with air. For all experiments, the airflow to the animal was held constant at 100 mL/min and odors were injected into a carrier stream. The absolute relative odor concentration was measured by a photoionization detector (PID; Aurora Scientific) and normalized to the largest detected signal for each odor. To create mixtures, air-phase dilution was used, and the total concentration of each odor was held constant. The measured mixture signal in the PID was nearly a perfect linear summation of the signal measured for each odor alone (Supplemental Figure 4). For all experiments, odors were delivered 2–6 times each.

For experiments characterizing the odor tuning of olfactory epithelium, the odor panel consisted of: 1) Ethyl tiglate 2) Allyl tiglage 3) Hexyl tiglate 4) Methyl tiglate 5) Isopropyl tiglate 6) Citronellyl tiglate 7) Benzyl tiglate 8) Phenylethyl tiglate 9) Ethyl tiglate 10) 2-Ethyl hexanal 11) Propyl acetate 12) 4-Allyl anisole 13) Ethyl valerate 14) Citronellal 15) Isobutyl proptionate 16) Allyl butyrate 17) Methyl propionate 18) Pentyl acetate 19) Valeric acid 20) (+)Carvone 21) (-)Carvone 22) 2-Methoxypyrazine 23) Isoeugenol 24) Butyl acetate 25) Valeraldehyde 26) Isoamyl acetate 27) Methyl valerate 28) Octanal 29) 2-Hexanone 30) Methyl butyrate 31) 2-Heptanone 32) Acetophenone. See supplemental Figure 4 for PID measurements. For experiments measuring complex mixture responses in the olfactory epithelium, odors 1-16 were used from the panel above.

### Data analysis

Images were processed using both custom and available MATLAB (Mathworks) scripts. Motion artifact compensation and denoising was done using NoRMcorre (Pnevmatikakis and Giovannucci, 2017). For experiments imaging OSN axon terminals in the olfactory bulb, regions of interest (ROIs) were manually selected by outlining glomeruli in maximum projection images. For epithelium imaging, the CaIMaN CNMF pipeline (Pnevmatikakis et al., 2016) was used to select and demix ROIs. ROIs were further filtered by size and shape to remove merged cells. For all mixture experiments, the peak ΔF/F signal was calculated by finding the peak signal following odor onset and averaging with the two adjacent points. The mean ΔF/F signal in the 20 frames following odor onset was used for odor tuning experiments. To account for changes in respiration and anesthesia depth, correlated variability was corrected for (Mathis et al., 2016). Thresholds for classifying responding ROIs were determined from a noise distribution of blank (no odor) trials from which three standard deviations were used for responses. Across all datasets, only ROIs with at least one significant odor response were included for further analysis. Measurements of binary mixture non-linearities used individual trial replicates of the three highest odor concentrations used in each experiment.

### Data fitting

We further analyzed the response curves from 344 glomeruli corresponding to Methyl tiglate, Isopropyl propionate, and their mixture, 226 glomeruli corresponding to Ethyl valerate, Allyl butyrate, and their mixture, and 964 ORNs corresponding to Methyl tiglate, Isopropyl propionate and their mixture. The observed Ca^2+^ fluorescence response *R(c)* was fit using a sigmoid function against log odor concentrations delivered over 3 orders of magnitude using the equation:

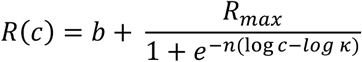

Here *b* sets the baseline response for blank odors, *R*_*max*_ + *κ* is the response at saturating concentrations and *κ*^−1^ is the binding affinity. We filtered for noisy data, unresponsive glomeruli/ORN-odor pairs and low sensitivity by imposing thresholds on the mean-squared error from the best fit parameters, the maximal response and high *κ* respectively.

## Acknowledgements

This work was partly supported by grants from the NIH (R01 DC014453 & DC016289) to VNM. JDZ was supported by NIH Fellowship F32 DC015938. GR was partially supported by the NSF-Simons Center for Mathematical & Statistical Analysis of Biology at Harvard (award number #1764269) and the Harvard Quantitative Biology Initiative. We also acknowledge support from the following grants, which facilitated critical discussions at KITP, Santa Barbara: NSF (PHY-1748958), NIH (R25 GM067110), and the Gordon and Betty Moore Foundation (2919.01)

## Author Contributions

JDZ, GR, MV, and VNM designed the experiments. JDZ. acquired the data. JDZ and GR analyzed the data. JDZ and VNM wrote the manuscript with inputs form all authors.

## Supplemental Figures

**Supplemental Figure 1.**
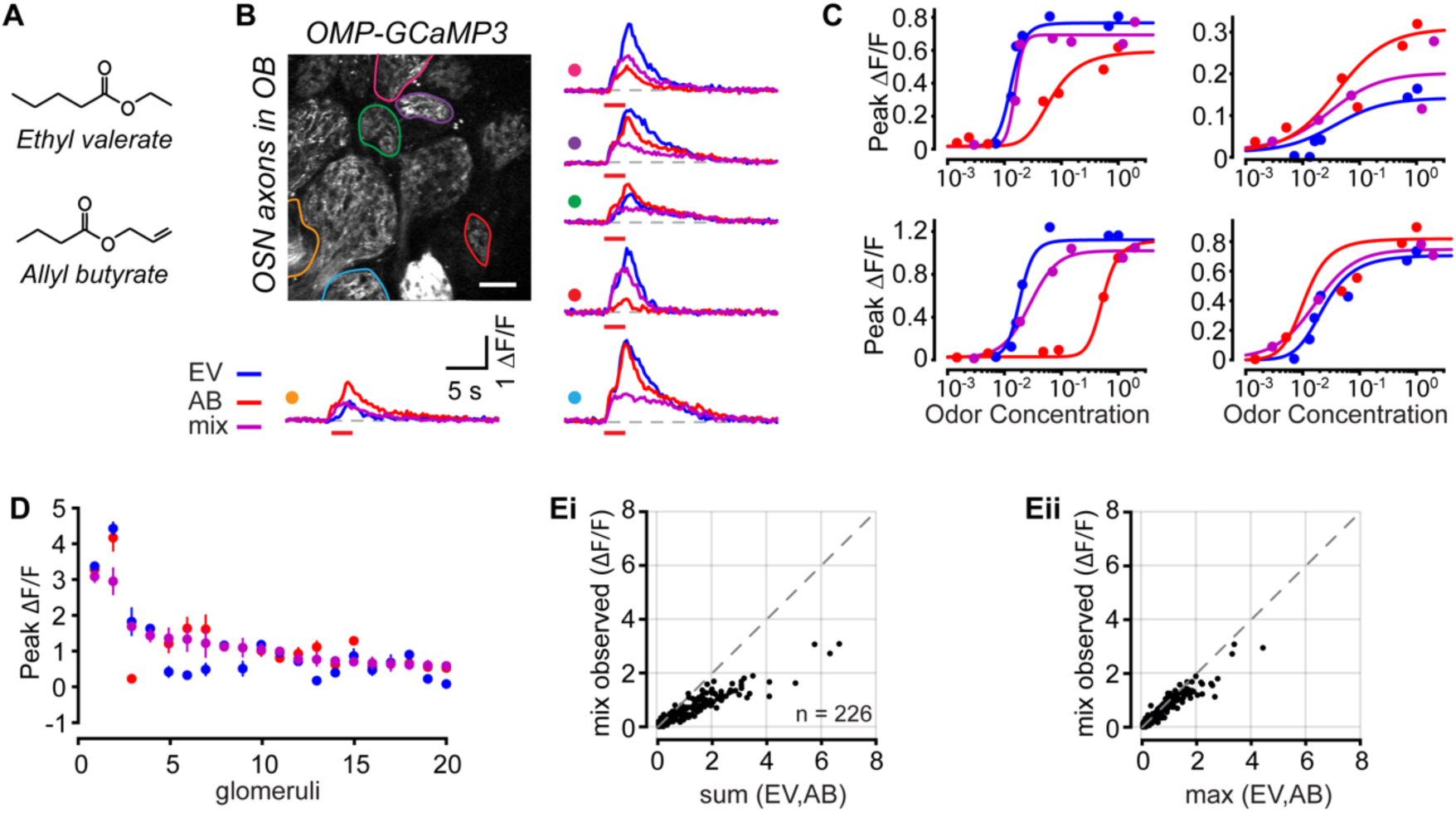
Antagonism measured in OSN axon terminals using a functionally overlapping odor pair. Related to Figure 1. **A**. Experimental setup. Odor responses in OSN axon terminals were measured at olfactory bulb glomeruli through a cranial window. **B**. Example image of glomeruli and selected ROIs. Odor responses from selected glomeruli are shown on right as ΔF/F time courses. Colored circles correspond to ROIs in image. Red bar under traces denotes odor delivery time. Scale bar in image is 50 µm. **C**. Example dose response curves from four selected glomeruli. Each point is the average of 3-5 trials. **D**. Data from 20 randomly selected glomeruli. Each point is the mean of the three largest responses from each odor. Error bars are s.e.m. **E**. (**i**) Comparison of the observed mixture response against the linear sum of both mixture components or (**ii**) against the maximum response generated by either mixture component.

**Supplemental Figure 2.**
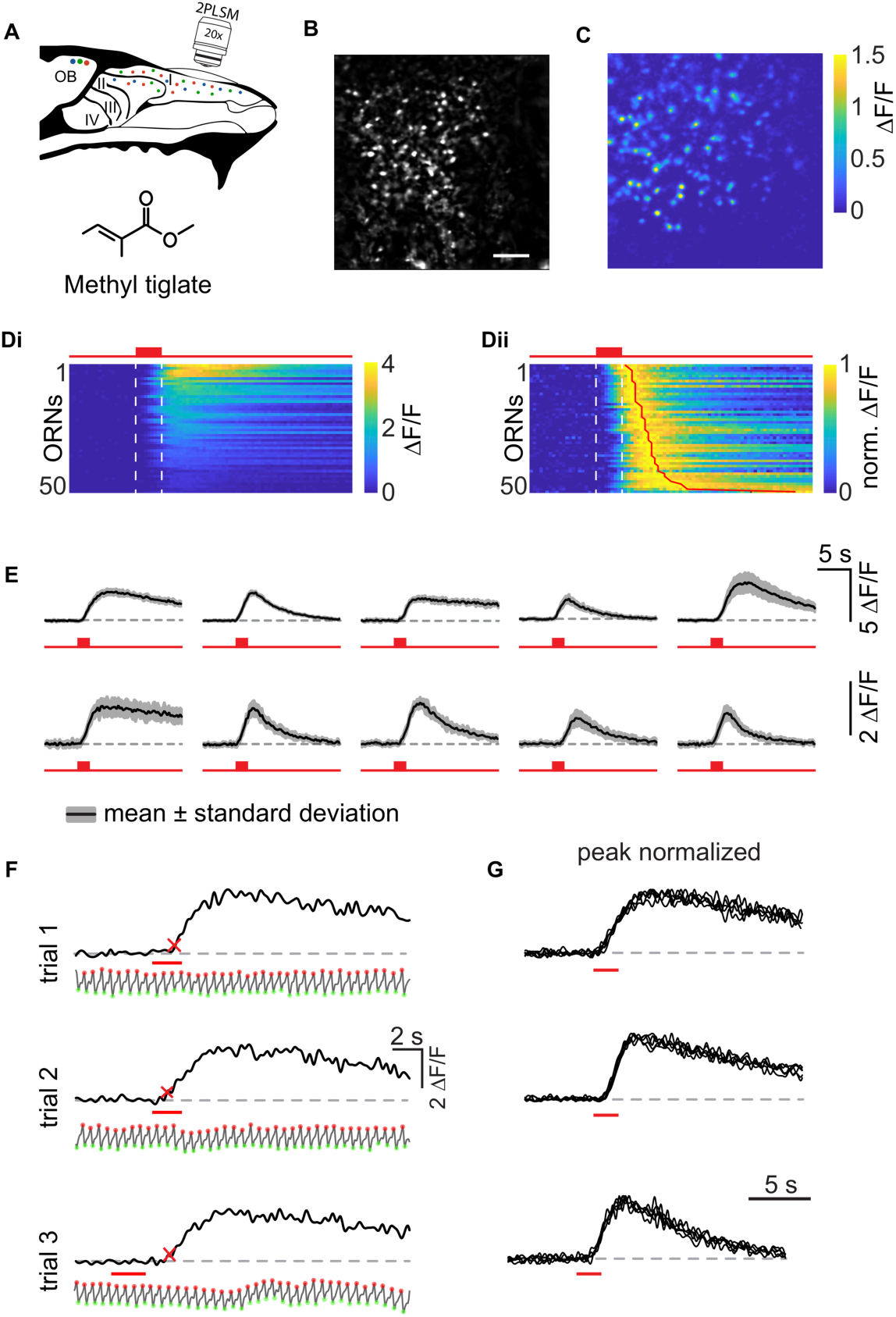
Response stability and kinetics of single OSNs. **A**. Experimental setup for imaging OSN somata in the epithelium for the odor Methyl tiglate. **B**. Resting fluorescence of OSN somata in the olfactory epithelium. Scale bar in image is 50 µm. **C**. ΔF/F image in response to the odor Methyl tiglate for the imaging field in B. **Di**. Time course of 50 selected OSNs sorted by response amplitude in response Methyl tiglate. **Dii**. Same 50 OSNs normalized to their peak amplitude and sorted by peak position. Red box and dashed lines indicate odor delivery time. **E**. 10 selected OSNs showing odor response time course. Shaded area is standard deviation from 10 repetitions of odor deliver. **F**. Three consecutive responses from the same OSN and accompanying respiration trace. Peak of inhalation and exhalation are denoted by red and green circles. Odor delivery time is red bar. Red cross denotes point when trace first deviates 3 S.D. from baseline. **G**. Three example OSNs in response to three odor repetitions. Top traces correspond to part *E*. Traces are normalized to their peak amplitude to show waveform consistency across trials for the same odor.

**Supplemental Figure 3.**
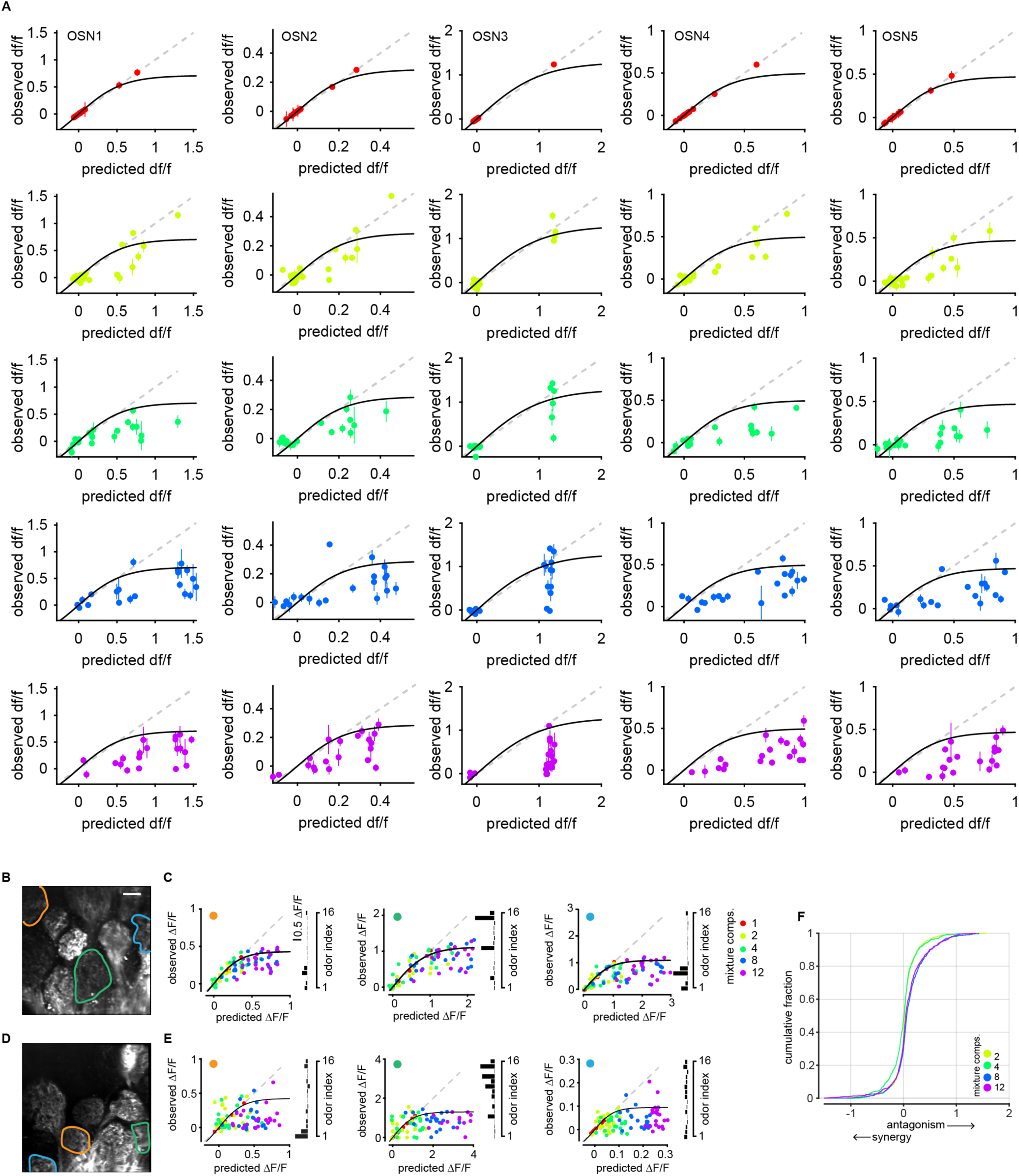
Mixture responses for single OSNs. Related to Figure 4. **A**. Mixture responses for five selected OSNs. Each column corresponds to a single OSN and rows are mixture complexities arranged by increasing mixture component number. Each point is the mean and s.e.m. of three repetitions of an odor mixture. Supplemental OSN1 corresponds to Figure 4 OSN2 and supplemental OSN3 corresponds to Figure 4 OSN4. **B**. Example image of glomeruli and selected ROIs. Scale bar is 50 µm. **C**. Example mixture responses from the glomeruli outlined in B. Each point is the average of three trials. The data from each glomerulus is fit with a sigmoid that reflects the maximum possible response. The odor tuning profile for each glomerulus is shown on the right. **D**. Additional example of glomeruli and selected ROIs from another animal. **E**. Same as part *B*. **F**. Cumulative distribution of all deviations from the sigmoidal fit for each mixture complexity. Data collected from 39 glomeruli in four mice.

**Supplemental Figure 4.**
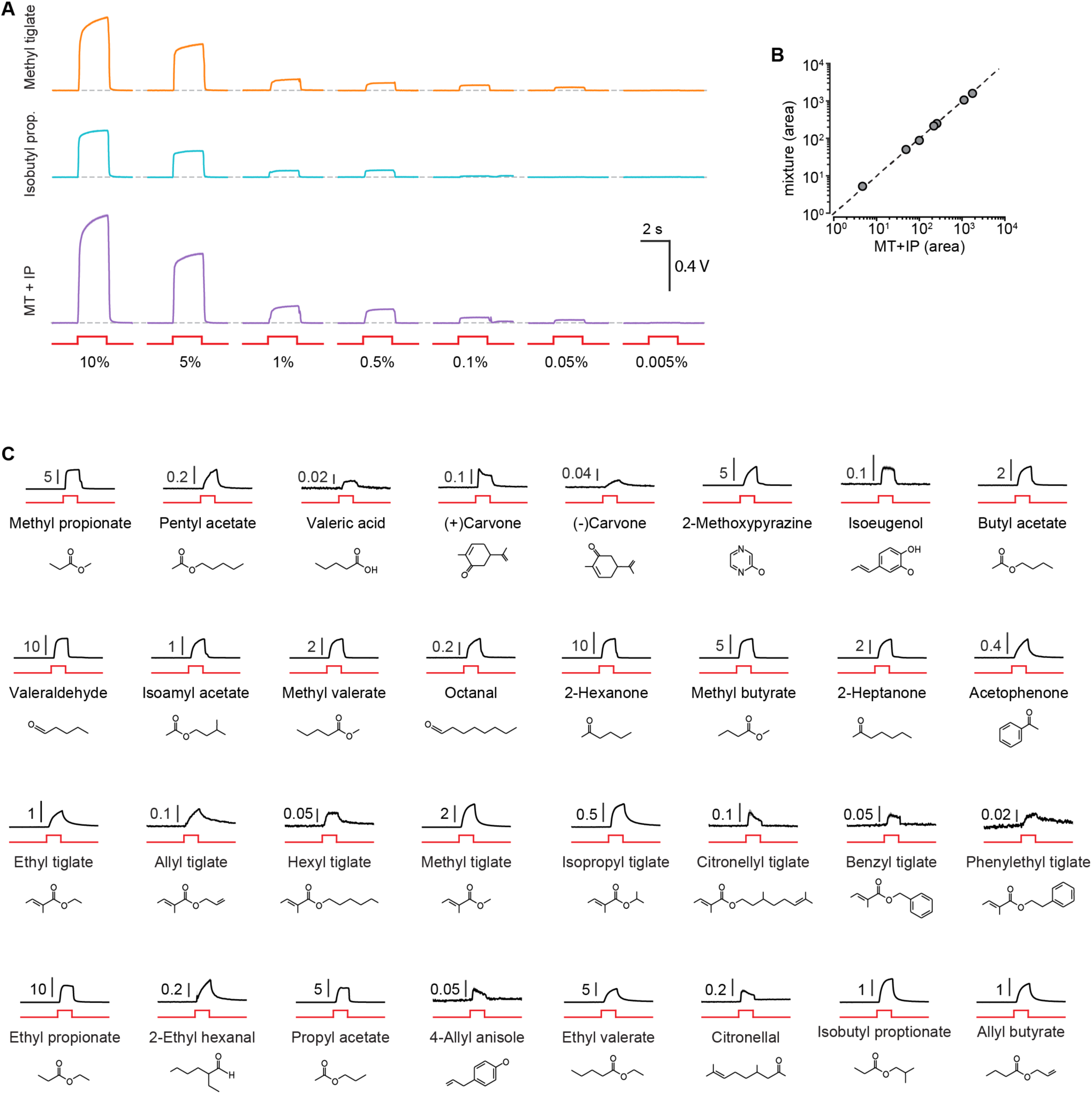
Photoionization detector (PID) traces for binary odor mixture experiments. **A**. PID traces for the odor pair Methyl tiglate and Isobutyl propionate at seven v/v dilutions. Solid lines are mean of five trials and shaded area is s.e.m. Odors were delivered for 2 s. Voltage command to the olfactometer is in red. Estimated concentration after air dilution is given for each trace below. **B**. For each v/v dilution the linear sum of the area under the PID trace for each odor component is compared to the measured PID measured mixture response. The mixture response is nearly a perfect summation of mixture components when delivered alone. **C**. PID traces of all odors used in tuning experiments in black (mean of 5 trials), voltage command to olfactometer in red. Molecular shape of each odor is below the corresponding traces.

